# Separate orexigenic hippocampal ensembles shape dietary choice by enhancing contextual memory and motivation

**DOI:** 10.1101/2023.10.09.561580

**Authors:** Mingxin Yang, Arashdeep Singh, Molly McDougle, Léa Décarie-Spain, Scott Kanoski, Guillaume de Lartigue

## Abstract

The hippocampus (HPC), traditionally known for its role in learning and memory, has emerged as a controller of food intake. While prior studies primarily associated the HPC with food intake inhibition, recent research suggests a critical role in appetitive processes. We hypothesized that orexigenic HPC neurons differentially respond to fats and/or sugars, potent natural reinforcers that contribute to obesity development. Results uncover previously-unrecognized, spatially-distinct neuronal ensembles within the dorsal HPC (dHPC) that are responsive to separate nutrient signals originating from the gut. Using activity-dependent genetic capture of nutrient-responsive HPC neurons, we demonstrate a causal role of both populations in promoting nutrient-specific preference through different mechanisms. Sugar-responsive neurons encode an appetitive spatial memory engram for meal location, whereas fat-responsive neurons selectively enhance the preference and motivation for fat intake. Collectively, these findings uncover a neural basis for the exquisite specificity in processing macronutrient signals from a meal that shape dietary choices.

## Introduction

Survival hinges upon the acquisition of sufficient food to meet metabolic demands. Therefore, possessing the capacity to construct a cognitive map and navigate accurately to a known food source within the environment confers a distinct competitive advantage. Animals learn to utilize contextual cues linked to the nutritional value of the food,^1^ and forming episodic memories of the spatial location of the cues enables efficient return to previously encountered food sources. Repeatedly associating discrete or contextual cues with food in a manner that predicts food intake induces a motivational state that amplifies the desire to eat - a phenomenon termed cue-potentiated eating.^2^ This adaptive behavior becomes overwhelmed in our current food environment characterized by an inundation of food-associated cues and readily-available foods rich in fats and sugars. Associative learning mechanisms linking food cues with intake of calorie-dense diets amplifies susceptibility to obesity development. Supporting this notion, brain reactivity to food cues predicts current weight status^3^, the inclination to gain weight in future^4, 5^, and food choice^6, 7^ Hence, unraveling mechanisms governing memory formation regarding contextual cues linked to fat and sugar intake holds potential for combating obesity.

The hippocampus (HPC) is a neural substrate critical for cognitive mapping^8^ and the formation of episodic memories related to autobiographical experiences and their contextual details^9, 10^. Given the pivotal role of navigational and contextual memory in acquiring food, it is not surprising that recent evidence suggests the HPC also plays a role in the control of food intake^11, 12^. Specifically, the HPC becomes activated by post-ingestive signals following a mixed meal,^13^ hormones released from the gut in response to eating,^13^ and sensory cues associated with meals, including odors,^14, 15^ taste,^16, 17^ texture,^18^ tones,^19^ and visual cues.^20^ HPC lesioning studies in rats have demonstrated an increase in food intake^21^ and body weight in both females^22^ and males.^11^ Conversely, chemogenetic stimulation of glutamatergic HPC neurons inhibits 24-hour food intake.^23^ Patients with retrograde amnesia resulting from brain lesions that encompass the HPC consume multiple successive meals,^24, 25^ which can be interpreted as impaired memory or impaired sensing of internal metabolic needs, an outcome also observed in rodents with HPC lesions.^26, 27^ Disruption of HPC function has also been associated with obesity. In a human fMRI study, hippocampal blood flow was lower after a meal in individuals that were obese compared to those of a healthy weight.^28^ Feeding rats a high-fat high-sugar diet impairs performance on hippocampal-dependent spatial learning and episodic memory tasks.^29^ Taken together, these data highlight the HPC as having an anorexigenic role in energy metabolism, with mechanisms involving episodic memory,^30, 31^ spatial memory,^32^ and appetitive reward.^26, 33^

The HPC has also been found to be activated in conditions associated with increased food intake. Ghrelin, an orexigenic hormone released from the stomach under fasting conditions,^34^ increases food intake and motivation to work for sugar reward when administered into the HPC of rats.^35^ In human fMRI studies, HPC activity is enhanced in response to images of food and tastants,^36, 37^ shown to promote arousal and motivation to eat,^38, 39^ and these effects are strongest in individuals with obesity. Recent findings identified an HPC subregion in humans as a key hub for encoding the appetitive value of sugar and fat, with a compromised HPC appetitive subnetwork in individuals with obesity.^40^ These data suggest a potential role for the HPC in increasing food intake, although the existence of a specific orexigenic population of HPC neurons remains unproven. This knowledge gap may partly stem from limitations in the temporal and spatial resolution of previous studies employing lesions and pharmacological approaches. Recent advances in transcriptomic analyses have unveiled extensive molecular diversity in HPC neurons^41–44^ and efforts continue to functionally characterize subpopulations of HPC neurons based on their projection patterns and/or genetic markers.^45^ Notably, screening of meal-responsive neurons revealed that a substantial number of neurons in the dHPC are activated by eating.^23^ Among these, a subset of neurons in the hilar region of the dHPC has been identified as expressing DRD2, and using molecular and genetic tools were demonstrated to inhibit food intake.^23^ These types of approaches provide an opportunity to identify orexigenic populations within the HPC. We hypothesized that fat and sugar may activate a subset of HPC neurons with orexigenic function. Our results find subsets of HPC neurons that are recruited in response to fats or sugars, and leverage Fos^TRAP^ mice as an unbiased approach to manipulate the activity of these HPC neurons to test their role in appetitive behavior.

## Results

### Dorsal hippocampal neurons are responsive to different post-ingestive nutrients

Previous studies demonstrate that the HPC is activated in response to mixed nutrient chow^23^ and following intragastric infusion of a mixed meal^13^. To test if the HPC is activated in response to individual reinforcing nutrients, we measured Fos immunofluorescence, a marker of neuronal activity, in mildly-fasted wildtype mice in response to intragastric infusions (500 µl, 100 µl/min) of sugar (sucrose, 15% w/v), equicaloric fat (microlipid, 6.8% v/v) or isosmotic saline (0.9% w/v) (Fig 1A). Fos was increased in discrete neuronal populations within the dorsal hippocampus (dHPC) in mice receiving infusions of sucrose or fat compared to saline (Fig 1B). Similar Fos density was found in response to both nutrients (Fig 1C), and the highest density of neurons were proportionally enriched in the dentate gyrus (DG) in response to both sucrose (Fig 1D) or fat (Fig 1E). Notably, intragastric infusions of sucrose and fat also resulted in similar density of Fos labeling in the ventral HPC (vHPC) (S1A-E), but in this region responsive neurons were particularly enriched in the CA1 (S1F-G). Together these data highlight that separate dHPC neurons are responsive to different post-ingestive nutrient signals from the gut.

**Figure 1.**
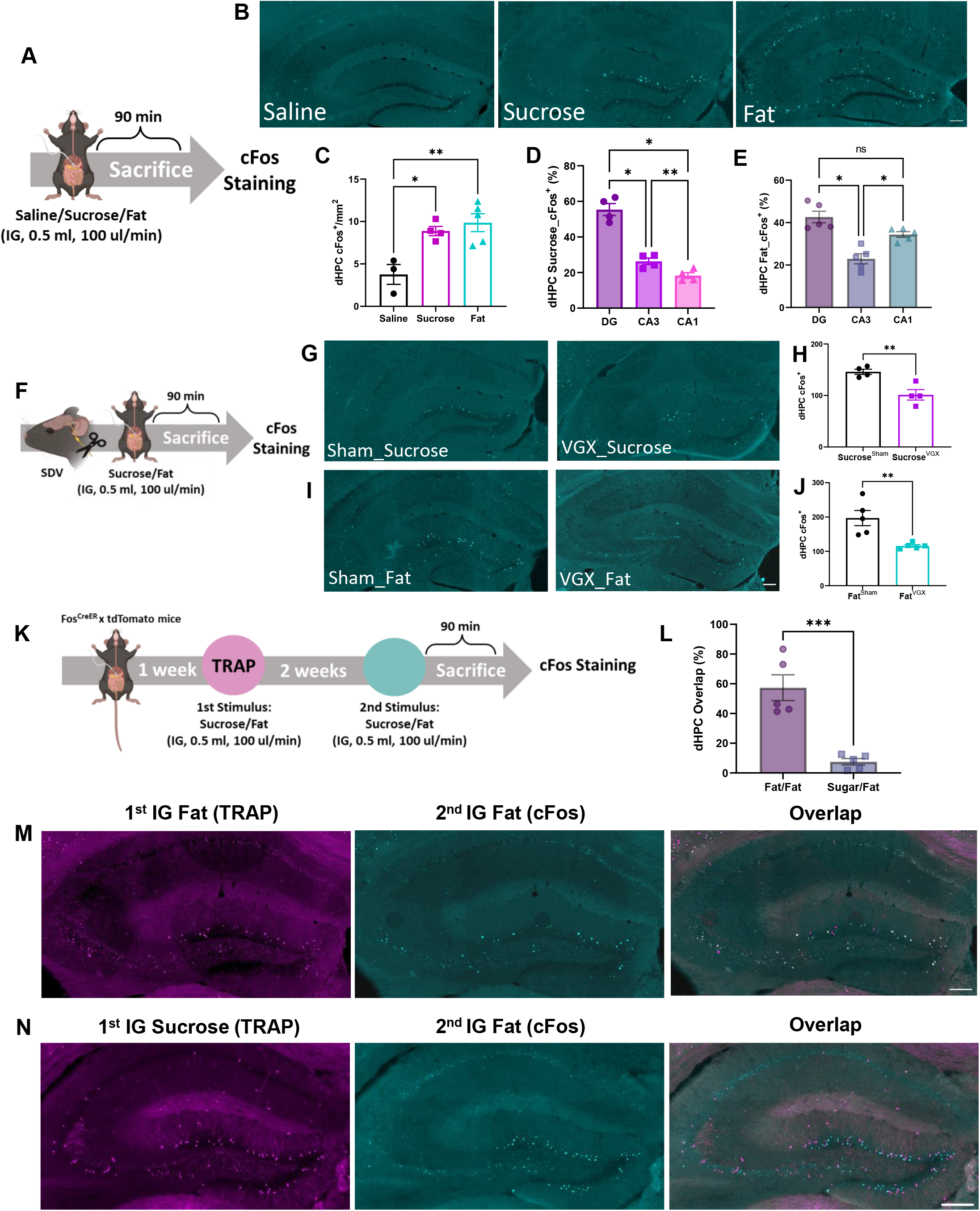
Dorsal hippocampal neurons are responsive to different post-ingestive nutrients. A Schematic of Fos staining approach to assess dHPC neuronal responsiveness to intragastric saline, sucrose (15%), or fat (6.8%). **B** Representative images of Fos expression in the dHPC. **C** Quantification of (B) showing increased Fos expression in dHPC neurons in response to sucrose or fat infusions compared to saline (N=3-5/group, one-way ANOVA with Turkey post hoc analysis). **D-E** Quantification of Fos expression in DG, CA3 and CA1 of the dHPC following intragastric sucrose (**D**) or fat (**E**) infusions (N=4-5/group, one-way ANOVA with Tukey post hoc analysis). **F** Schematic illustration of SDV for evaluating the role of the vagus nerve in dHPC neuronal nutrient sensing. **G** Representative images of Fos expression in the dHPC following intragastric sucrose infusion in mice with or without VGX. **H** Quantification demonstrating dHPC Fos expression in response to sucrose is blunted by VGX (N=4/group, unpaired t test). **I** Representative images of Fos expression in the dHPC following intragastric fat infusion in mice with or without VGX. **J** Quantification demonstrating dHPC Fos expression in response to fat is blunted by VGX (N=5/group, unpaired t test). **K** Schematic of Fos^TRAP^ approach comparing tdTomato labeling to Fos labeling in response to intragastric nutrient infusions. **L** Quantification showing higher overlap between repeated infusions of fat compared to separate macronutrients in the dHPC (N=5/group, unpaired t test). **M-N** Representative images of the dHPC in response to Fat (TRAP, magenta), and (**M**) colocalization after infusion two weeks later of Fat (Fos, cyan; top) or (**N**) fat (Fos, cyan; bottom) in the same animal. Data are presented as mean ± s.e.m. *P < 0.05, **P < 0.01, ***P < 0.001, NS, not significant. Scale bars 100 µm.

The vagus nerve is a key neural pathway that connects the gut and the brain. Subsets of vagal sensory neurons in the nodose ganglia (NG) are activated in response to intestinal nutrients, and these NG neurons are necessary to mediate the reinforcing value of fat and sugar.^46^ Although vagal sensory fibers terminate in the nucleus tractus solitarius (NTS) of the hindbrain, there is evidence of a polysynaptic circuit connecting the gut via the NTS to the HPC.^47^ Furthermore, vagal stimulation increases HPC activity in mice^48^ and in humans^49^ while deletion of gut-innervating vagal sensory neurons impairs HPC-dependent contextual episodic memory.^47^ ^50^ To test whether post-ingestive fats and/or sugars require the vagus nerve to recruit dHPC neurons, we quantified Fos expression in the dHPC in response to intragastric nutrient infusions in mice following subdiaphragmatic vagotomy (SDV) or sham surgery (Fig 1F). Nutrient-induced dHPC Fos expression was impaired in SDV animals, significantly reducing response to intragastric sucrose (Fig 1G-H) and fat (Fig 1I-J). These data demonstrate the vagus nerve acts as an important neural relay connecting nutrient signals in the gut to the dHPC.

We recently reported that separate NG populations sense fat or sugar and that these nutrients activate separable downstream central circuits.^46^ Thus, we inquired whether separate populations of dHPC neurons are recruited in response to fat and sugar. We used a Fos^TRAP^ mouse^51^ with a previously validated approach^46^ to compare neuronal activity in response to two separate nutrient infusions in the same mouse (Fig 1K). These mice express an inducible Cre recombinase, iCreER^T2^, under the control of an activity-dependent Fos promoter (*Fos^TRAP^* mice), enabling permanent genetic access to neuronal populations based on their activation to a specific, time-restricted stimulus.^52–54^ To validate the approach in the HPC, we compared the number of Fos^TRAP^ positive neurons and Fos immunofluorescence and found that the density of responsive neurons within the dHPC was similar (S1I-J). When analyzing the overlap between the repeated infusion of the same stimulus within the same animal, we found a high level of overlap in the dHPC between Fat^TRAP^ tdTomato labeling and Fat Fos immunofluorescence (Fig 1L-M). However, when comparing the response to different stimuli, there was low overlap in the dHPC neurons between tdTomato labeling of the Sucrose^TRAP^ neurons and Fos labeling of fat responsive neurons (Fig 1L, N). The difference between overlap in Fat^TRAP^/Fat^cFos^ mice and Sugar^TRAP^/Fat^cFos^ was particularly large in the dDG (S1H), consistent with the role of the DG as a pattern segregator^50–53^ and suggesting that dDG neurons may play a role in differentiating responses to different types of interoceptive information from the gut. Interestingly, we also find that IG infusion of starch (cornstarch, 15% w/v) colocalizes with dHPC neurons that are trapped with equicaloric solution of sucrose (15% w/v, S1K-L), suggesting that these neurons are broadly tuned to carbohydrates. Importantly, neither fat nor sucrose infusion resulted in activation of neurons of the hilar region of the dHPC (Fig 1B), suggesting that these populations are distinct from the previously described DRD2-expressing hilar neurons of the dHPC known to inhibit food intake.^23^ Thus, we identify two new populations of dHPC neurons that are responsive to separate post-ingestive nutrients that both involve vagally-dependent signaling mechanisms.

### Fat- and sugar-responsive dHPC neurons control nutrient-specific preference and intake

Next, we wanted to determine the role of nutrient-responsive dHPC populations in the control of food intake, and reasoned that these spatially segregated populations recruited by separate post-ingestive nutrients may differentially resolve food intake at the macronutrient level.^55, 56^ To genetically access dHPC neurons that were active in response to intragastric infusion of fat or sugar, we used TRAP2 mice (Fig 2A).^57, 58^ To assess necessity of these neurons in the control of feeding behavior, we injected a cre-dependent virus expressing caspase in the construct AAV-flex-taCasp3-TEVp.^59^ This approach allowed selective lesioning of dHPC neurons that respond to either fat (6.8%) or sugar (15%) compared to a control mouse that received dHPC viral injection that did not cause lesioning. Caspase treatment ablated sugar-responsive and fat-responsive dHPC neurons, as demonstrated by the greater than 50% loss in tdTomato-positive neurons in the caspase-treated mice compared to controls (Fig2B-D). To behaviorally assess the role of dHPC neurons in nutrient preference, we presented the mice with a choice between two bottles containing equicaloric solutions of either fat (6.8%) or sugar (15%) and quantified intake using lickometers (Fig 2A). Over three test days the control mice exhibited a preference for the fat solution over the sucrose solution (Fig 2E-H). The mice with ablated sugar-responsive dHPC neurons significantly decreased sucrose consumption compared to controls (Fig 2E), with no effect on fat intake (Fig 2F). Deletion of fat-responsive dHPC neurons resulted in no change in sucrose intake (Fig 2G), but reduced fat consumption by 40% compared to control mice (Fig 2H). The reduction in nutrient intake does not appear to be in response to reduced taste^60^ since there was no group differences in lick numbers over the first 10 seconds for either fat or sucrose between groups (S2A-B). Furthermore, deletion of sucrose-responsive dHPC neurons had no effect on sucrose or fat intake in a one bottle task, suggesting a primary role of dHPC^Sugar^ neurons in sucrose preference (S2C-D). Ablation of dHPC^Fat^ neurons had no effect on sucrose intake in a one bottle task (S2E), but reduced the number of licks for fat (S2F). In summary, these data demonstrate that separate populations of dHPC neurons are necessary for nutrient-specific preference.

**Figure 2.**
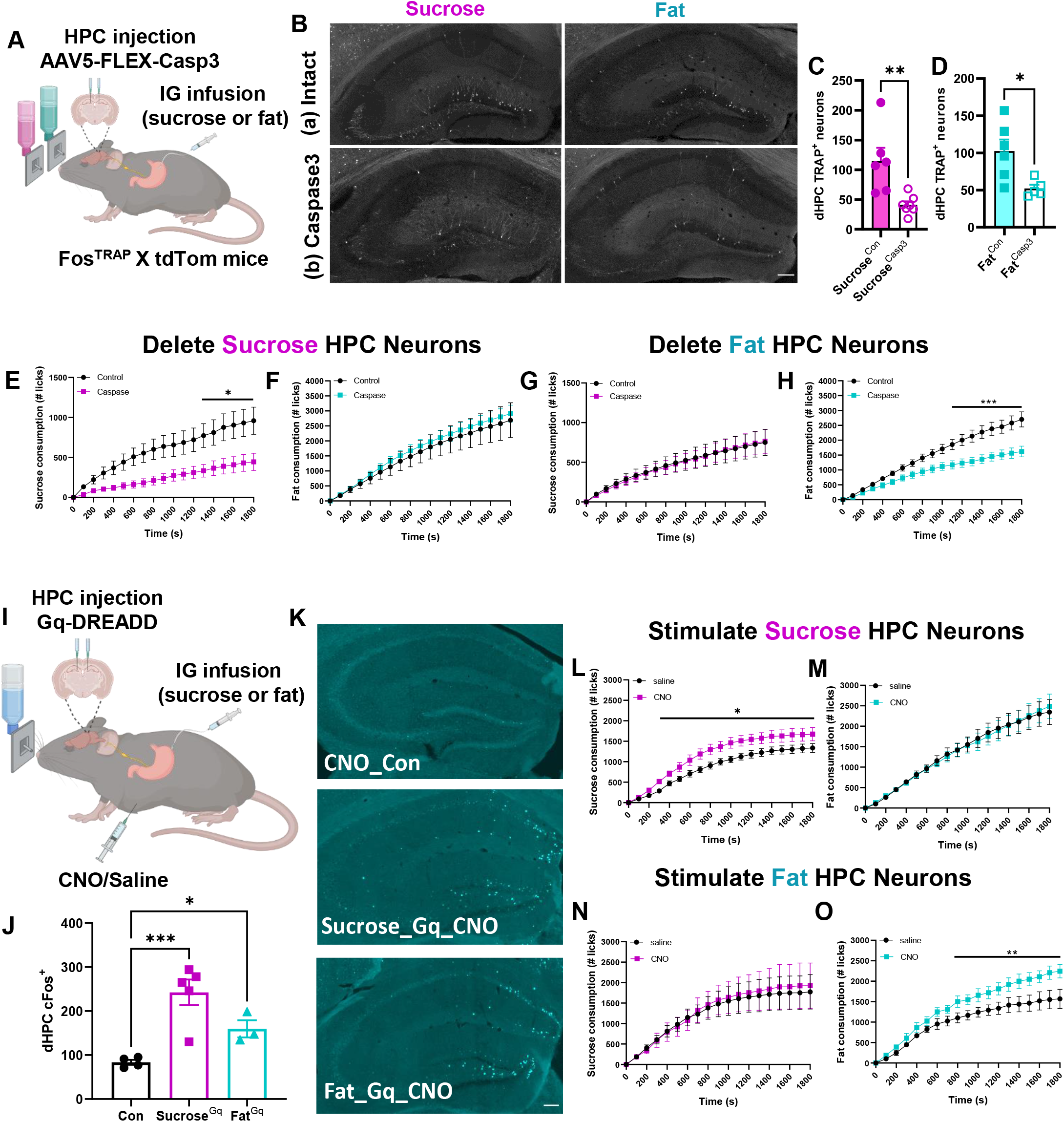
Fat- and sucrose-responsive dHPC neurons control nutrient-specific preference. **A** Schematic of the Fos^TRAP^ approach to selectively ablate nutrient-responsive HPC neurons that respond to intragastric infusion of sucrose or fat. **B** Representative images of nutrient-responsive HPC neurons from Fos^TRAP^ mice following intragastric infusion of sucrose or fat with or without viral-mediated caspase ablation. **C-D** Quantification demonstrating caspase deletion of tdTomato neurons (N=5-7/group, unpaired t test) in response to (**C**) sucrose or (**D**) fat. **E** Intake in a two-bottle choice test in mice with ablation of dHPC^Sugar^ neurons reduces sucrose solution consumption, **F** without affecting fat solution intake when sucrose and fat solution are both presented (N= 8-9/group, two-way ANOVA with Holm-Sidak post hoc analysis). **G** Intake in a two-bottle choice test in mice with ablation of dHPC^Fat^ neurons has no effect on sucrose intake, but **H** reduces fat solution consumption without affecting sugar solution intake (N= 8/group, two-way ANOVA with Holm-Sidak post hoc analysis). **I** Schematic of the Fos^TRAP^ approach to selectively stimulate nutrient-responsive HPC neurons that respond to intragastric infusion of sucrose or fat. **J** Quantification injection to demonstrated increased Fos labeling in dHPC after CNO in dHPC^Fat^ and dHPC^Fat^ mice expressing Gq-DREADD compared to control mice (N=4-5/group, one-way ANOVA with Two-stage linear step-up procedure of Benjamini, Krieger and Yekutieli post hoc analysis). **K** Representative images of Fos labeling in the dHPC after CNO. **L** Stimulation of sucrose-responsive dHPC neurons increases sucrose consumption **M** without affecting fat consumption (N=4-5/group, two-way ANOVA with Holm-Sidak post hoc analysis). **N** Stimulation of fat-responsive dHPC neurons has no effect on sucrose consumption, but **O** increases fat consumption without affecting sucrose consumption (N=4/group, two-way ANOVA with Holm-Sidak post hoc analysis). Data are presented as mean ± s.e.m. *P < 0.05, **P < 0.01, ***P < 0.001, NS, not significant. Scale bars 100 µm.

To assess the sufficiency of dHPC neurons in macronutrient preference, we performed chemogenetic stimulation of fat- or sugar-responsive dHPC neurons. A Cre-inducible viral Gq-coupled designer receptor encoded in the construct AAV-EF1a-DIO-hM3Dq-mCherry^61^, were bilaterally injected into the dHPC of Fos^TRAP^ mice (Fig 2I). CNO injection increased Fos expression in dHPC neurons in both Sugar^TRAP^ and Fat^TRAP^ mice expressing hM3Dq (Fig 2J-K), confirming our ability to chemogenetically activate Fos^TRAP^ HPC neurons in a nutrient-specific manner. In the one bottle task described above, chemogenetic activation of dHPC^Sugar^ neurons increased sucrose intake compared to vehicle treatment (Fig 2L), but had no effect on fat intake (Fig 2M). Stimulation of dHPC^Fat^ neurons exclusively increased fat consumption (Fig 2N-O). Importantly, none of these effects were observed when CNO was injected in mice not carrying the chemogenetic construct (S2G-H). These data suggest that the dHPC is attuned to specific macronutrients allowing for highly-refined feeding decisions.

### Fat- and sugar-responsive dHPC neurons control nutrient-specific episodic spatial memory

Next, we wanted to address the mechanisms by which dHPC neurons control nutrient-specific intake. The HPC forms context-specific neural representations that provide a physiological substrate of spatial memory^62^, and HPC activity is altered by contextual features of rewarding stimuli.^63–65^ To address whether dHPC^Sugar^ and dHPC^Fat^ neurons retain contextual information about the location of natural reinforcers, such as post-ingestive fats and sugars, we adapted a previously described food cup task.^23^ Mice were habituated to a novel context with two empty petri dishes, and during the training phase, one petri dish contained droplets of water while the other contained droplets of fat (6.8% v/v) or sucrose (15% w/v) solutions (Fig 3A). After training to learn the location of a nutrient-containing dish, we tested the mice with empty petri dishes in the same context to determine if they could remember the location of the nutrient-paired quadrant (Fig 3A). Control mice discriminated the sugar-paired quadrant above chance in tests 1h and 24h after the final training session (Fig 3B), suggesting that they were able to learn and remember the location of sucrose. Mice with ablated dHPC^Sugar^ neurons failed to discriminate the location of the sugar dish in the 1h and 24h tests (Fig 2C). However, when these mice repeated the task with fat solution using different contextual cues, both the control and sugar-ablated mice spent more time exploring the fat location at both timepoints (Fig 3D-E). In a separate group of mice trapped with fat, we found that controls and dHPC^Fat^ ablated mice were able to discriminate the sugar location in both 1 and 24h tests compared to the pretest (Fig 3F-G). Although the control mice exhibited fat location memory (Fig 3H), the ablation of fat-responsive dHPC neurons abolished the ability to discriminate the fat-paired location in both 1h and 24h tests (Fig 3I). Importantly, the order in which the nutrients were presented was counterbalanced, and there was no residual preference for the previous nutrient location following a 7-day washout period (S3A), favoring exploration of the new petri dish locations.

**Figure 3.**
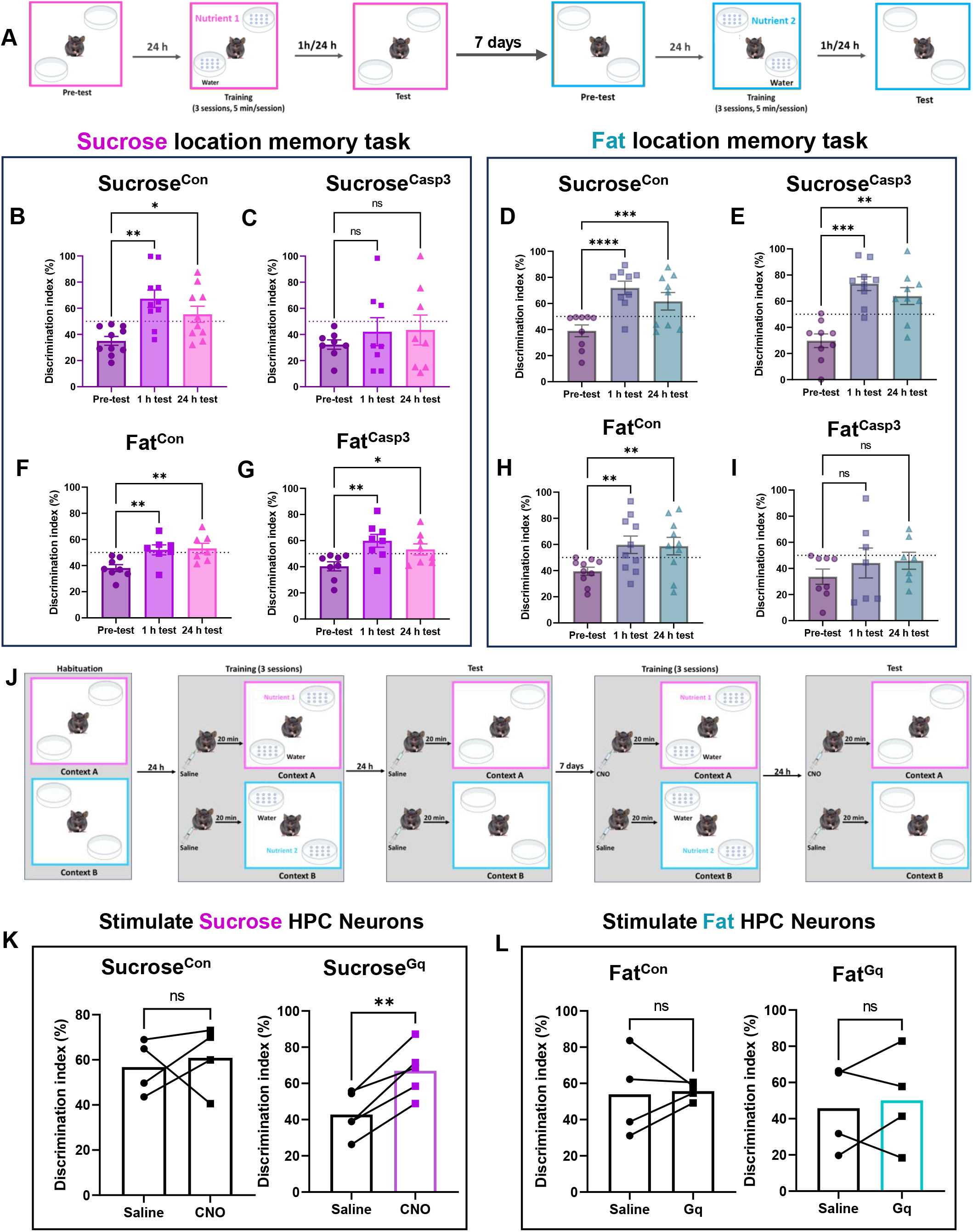
Fat- and sucrose-responsive dHPC neurons control nutrient-specific episodic spatial memory. **A** Schematic of nutrient-driven location memory task to assess the necessity of nutrient-responsive dHPC neurons in food-related reference memory. **B** Control mice showing increased discrimination for sucrose-paired location. **C** Ablation of dHPC^Sugar^ neurons prevents sugar location memory. **D** Control mice and **E** caspase-treated dHPC^Sugar^ mice form fat-driven memory (N=8-10/group, one-way ANOVA with Holm-Sidak post hoc analysis). **F** Fat control mice, and **G** mice with ablated dHPC^Fat^ neurons form sucrose memory. **H** Control mice spend more time exploring fat location after training, while **I** fat-ablated mice do not form location memory for fat. (N=7-10/group, one-way ANOVA with Holm-Sidak post hoc analysis). **J** Schematic of nutrient-driven location memory task to assess whether stimulation of nutrient-responsive dHPC neurons can improve nutrient-related memory. **K** Stimulation of dHPC^Sugar^ neurons improves sucrose-related memory recall (N=4-5/group, paired Student’s t test). **L** Stimulation of dHPC^Fat^ neurons does not improve fat-related memory recall (N=4/group, paired Student’s test). Data are presented as mean ± s.e.m. *P < 0.05, **P < 0.01, ***P < 0.001, NS, not significant.

To confirm that generalized spatial memory is not impaired, we performed a hippocampal-dependent^66^ novel object in context (NOIC) task (S3B). As expected, control mice spent more time exploring the object that is novel to the context, and similarly the ablation of either fat- or sugar-responsive dHPC neurons had no effect on the time spent exploring the novel object (S3C-F). These data indicate that the loss of nutrient-responsive dHPC neurons influences contextual memory of nutrient location, but that these neurons are specific to food and do not impair contextual memory for non-food related objects.

Increasing evidence suggests that the HPC is involved in working memory,^67, 68^ that allows retention of a small amount of information for a short period of time. To assess whether the nutrient-responsive dHPC neurons influence working memory related to food location, we performed a modified Barnes maze task.^47^ Mice were positioned in the center of a circular table and 8 petri dishes containing water solution and one containing either sucrose (15% w/v) or equicaloric fat (6.8% v/v) were evenly distributed around the edge (S3G). The location of the nutrient-containing dish remained the same across two consecutive trials per day, but changed each subsequent day. The index of working memory on this task is the difference in the number of errors (exploration of water dishes) between trials on 3 individual experimental days. We observed no difference in the number of errors between any of the groups in response to sugar (S3H-K) or fat (S3L-O), suggesting that ablation of nutrient-responsive dHPC neurons does not play a role in working memory. Altogether, these data suggest that both fat- and sugar-responsive dHPC neurons are necessary for episodic spatial memory for the location of individual nutrients.

Next, we assessed whether activation of nutrient-responsive dHPC neurons can improve context-dependent spatial memory for individual nutrients. Mice expressing hM3Dq or control virus in dHPC neurons trapped with intragastric infusion of either sucrose (15%) or fat (6.8%) were habituated to two novel contexts. During a 3-day training phase the mice receive saline injections 20 minutes prior to being placed in context A in the morning to learn to associate the location of a nutrient-containing dish for 10 min, and then received another saline injection before being placed in context B in the afternoon to learn a different location for the second nutrient-containing dish. Twenty-four hours later the mice were tested to determine if they could discriminate the correct context-specific nutrient-paired quadrant (Fig 3J). The mice failed to discriminate context-specific locations of sucrose or fat when treated with saline (Fig 3K-L). After 7 days the same test was repeated but the mice received CNO (3 mg/kg, IP) during training and test days before they were reintroduced into the context that had been previously paired with the specific nutrient with which they were initially trapped. To avoid desensitization, CNO was not injected on the third training day. We found that CNO had no impact on the performance of control mice (Fig 3K-L); however, chemogenetic stimulation improved the discrimination of hM3Dq-expressing dHPC^Sugar^ mice in response to CNO compared to vehicle treatment (Fig 3K). No improvement was observed in response to chemogenetic stimulation of dHPC^Fat^ neurons (Fig 3L). These data indicate that sugar-responsive dHPC neurons encode an engram of spatial and context-dependent memory for sugar.

### Fat-responsive HPC neurons encode motivation for fat

Dietary preferences are largely learned^69^ and this process involves reward-based associations.^46, 52, 69, 70^ In light of the increased preference for nutrients caused by dHPC neurons (Fig 2), we hypothesize that these neurons are involved in reinforcement learning. To address this, we assessed the role of nutrient-responsive dHPC neurons in a flavor-nutrient conditioning task, in which animals are trained to prefer a novel non-nutritive flavor that has been experimentally paired to an intragastric infusion of nutrient (Fig 4A).^71^ During conditioning, both control and caspase-treated mice with ablated dHPC^Sugar^ neurons received the same number of sucrose (15%) infusions (S4A), suggesting that sucrose resulted in similar levels of appetition in both groups.^72^ After conditioning, both control mice and caspase-treated mice formed increased preferences for the flavor paired with intragastric infusion of sucrose compared to their initial flavor preference (Fig 4B). Notably, control mice retain the flavor preference (Fig 4C), while the caspase-treated mice forget the conditioned preference by day 3 (Fig 4D). In mice lacking dHPC^Fat^ neurons the number of conditioning infusions of intragastric fat (6.8%) were severely reduced compared to control mice (S4B). After conditioning, control mice increased preference for the flavor paired with fat, while loss of dHPC^Fat^ neurons prevented fat reinforcement learning (Fig 4E). These data highlight different functions of the separate populations of nutrient-responsive dHPC neurons between formation and retention of conditioned preferences.

**Figure 4.**
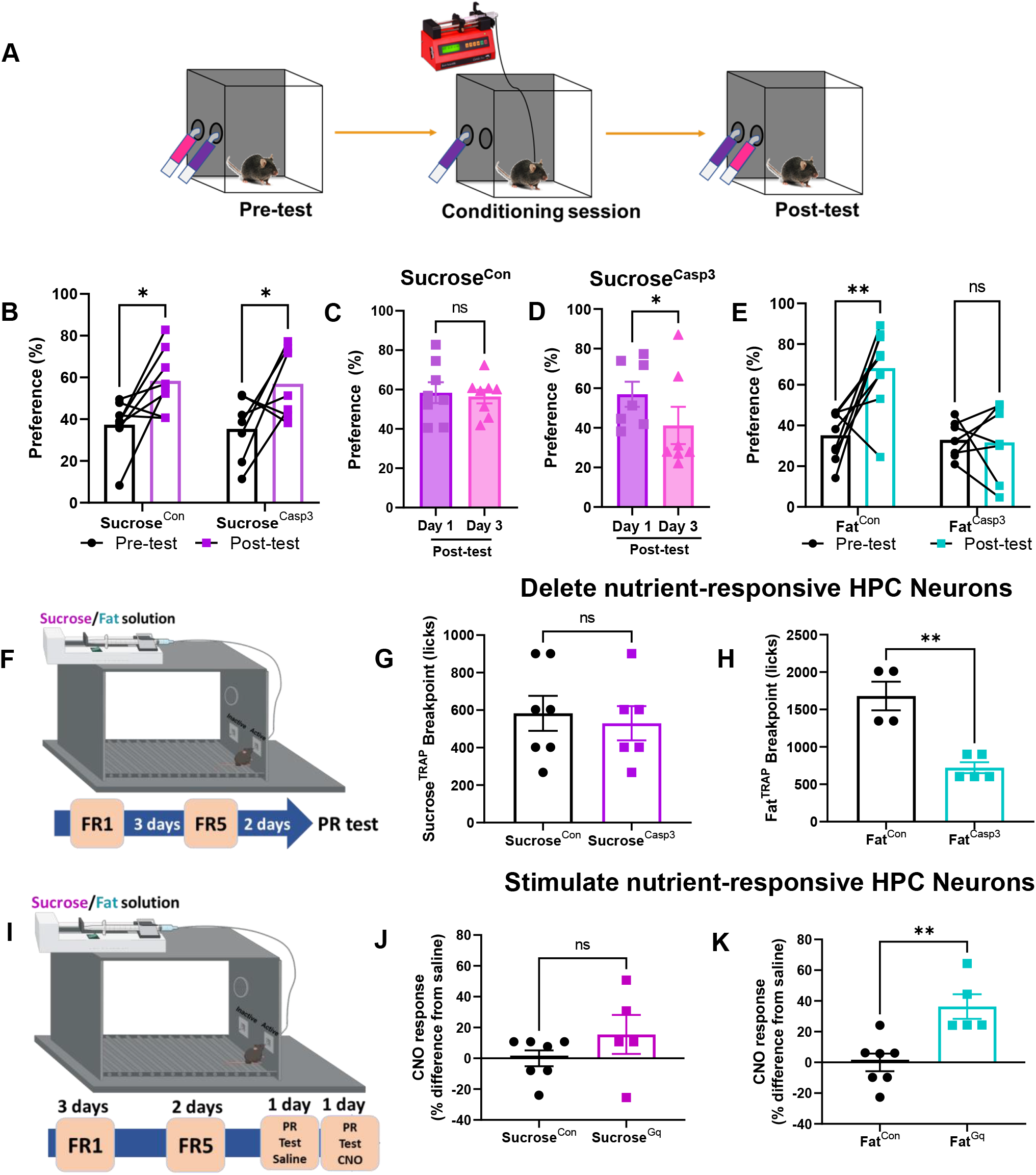
Fat-responsive dHPC neurons promote fat reinforcement. **A** Diagram demonstrating the flavor-nutrient conditioning paradigm. **B** Conditioning increases preference for the flavor paired with intragastric sucrose in Sugar^TRAP^ mice (N=7-8/group, two-way ANOVA with Holm-Sidak post hoc analysis). **C** Control mice remember the flavor preference 3 days post-training, while **D** caspase-treated dHPC^Sugar^ mice reduce preference by day 3. (N=7- 8/group, paired Student’s t test). **E** Ablation of fat-responsive dHPC neurons prevents fat reinforcement (N=7-9/group, two-way ANOVA with Holm Sidak post hoc analysis). **F** Schematic illustration of progressive ratio licking test to assess necessity of nutrient-responsive dHPC in motivation. **G** Ablation of sucrose-responsive dHPC neurons has no effect on sucrose motivation (N=6-7/group, unpaired t test). **H** Ablation of fat-responsive dHPC neurons reduces fat motivation, indicated by the decrease in breakpoint (N=4-5/group, unpaired t test). **I** Schematic illustration of progressive ratio licking test assessing sufficiency of nutrient-responsive dHPC neurons in motivation. **J** Stimulation of dHPC^Sugar^ neurons has no effect on sucrose motivation (N=5-7/group, unpaired t test). **K** Stimulation of Gq DREADD expressing dHPC^Fat^ neurons promotes fat motivation in response to CNO compared to the baseline (N=5-7/group, unpaired t test). Data are presented as mean ± s.e.m. *P < 0.05, **P < 0.01, ***P < 0.001, NS, not significant.

There is evidence that the dHPC is involved in motivation,^73^ thus we next assessed if nutrient-responsive dHPC neurons will increase the motivation for food. Effort-related motivation can be assessed by testing behavior using progressive ratio (PR) schedule reinforcement.^74^ We used an exponential PR task to probe the willingness of mice to lick for a dry sipper that requires an increasing number of licks for a small nutrient reward (Fig 4F). We quantified the number of licks required before an animal ceases to be willing to expend effort for a single reward, known as the breakpoint.^75^ Deletion of dHPC^Sugar^ neurons had no impact on the willingness to work for sucrose compared to control mice (Fig 4G). However, deletion of dHPC^Fat^ neurons reduced the breakpoint for fat compared to control mice, suggesting an important role of dHPC^Fat^ neurons in motivation (Fig 4H). During the training phase, we observed no group differences in the discrimination for the active nose hole or the number of licks the animals performed under FR1 or FR5 ratio (S4C-D). Next, we tested whether stimulation of dHPC neurons could increase the motivation to work for nutrients using a similar PR task as above (Fig 4I). All mice rapidly learned to discriminate the active nose hole to receive a small nutrient droplet triggered by a pump under FR1 schedule and FR5 schedule (S4E-F). After training, the mice were tested on an exponential PR schedule in response to saline or CNO on subsequent days. CNO (3 mg/kg, IP) had no effect compared to saline on sucrose breakpoint in dHPC^Sugar^ control mice or hM3Dq mice (Fig 4J). CNO significantly increased the willingness to nose poke for a small fat reward in the mice expressing hM3Dq, but had no effect in mice that did not express the chemogenetic construct (Fig 4K). Together these data identify a novel population of neurons in the dHPC that are necessary and sufficient for the motivation to consume fat.

### Nutrient-sensing dHPC neurons guide food intake based on diet composition

Having demonstrated the importance of dHPC neurons in memory, preference and motivation for individual macronutrients, we next wanted to assess the necessity of these neurons in the control of consumption of complex diets with mixed nutrient composition. We monitored continuous and uninterrupted ad libitum chow intake. Caspase-treated mice with ablated dHPC^Sugar^ neurons ate significantly less over 24 hours compared to control mice (Fig 5A). The hypophagia in the caspase-treated mice was caused by smaller meal size compared to controls, with no effect on meal duration or frequency (Fig 5B). We also compared food intake within animals before and after caspase-mediated neuron ablation, and found that the caspase-treated mice ate significantly less post-compared to pre-ablation (S5A), but no difference in food intake was observed in the control mice before and after TRAP (S5B). There were also no group differences in food intake pre-ablation (S5C). Deletion of dHPC^Fat^ neurons had no effect on chow intake (Fig 5C) or meal patterning (Fig 5D) compared to controls. There were no within animal differences before or after undergoing the fat TRAP protocol (S5D-F). These data indicate subpopulations of dHPC neurons increase daily cumulative food intake. The fact that only dHPC^Sugar^ neurons increased intake of chow, a carbohydrate-rich diet, suggests that dHPC neurons are attuned to select nutrients in a mixed meal and selectively increase food intake according to the composition of the diet.

**Figure 5.**
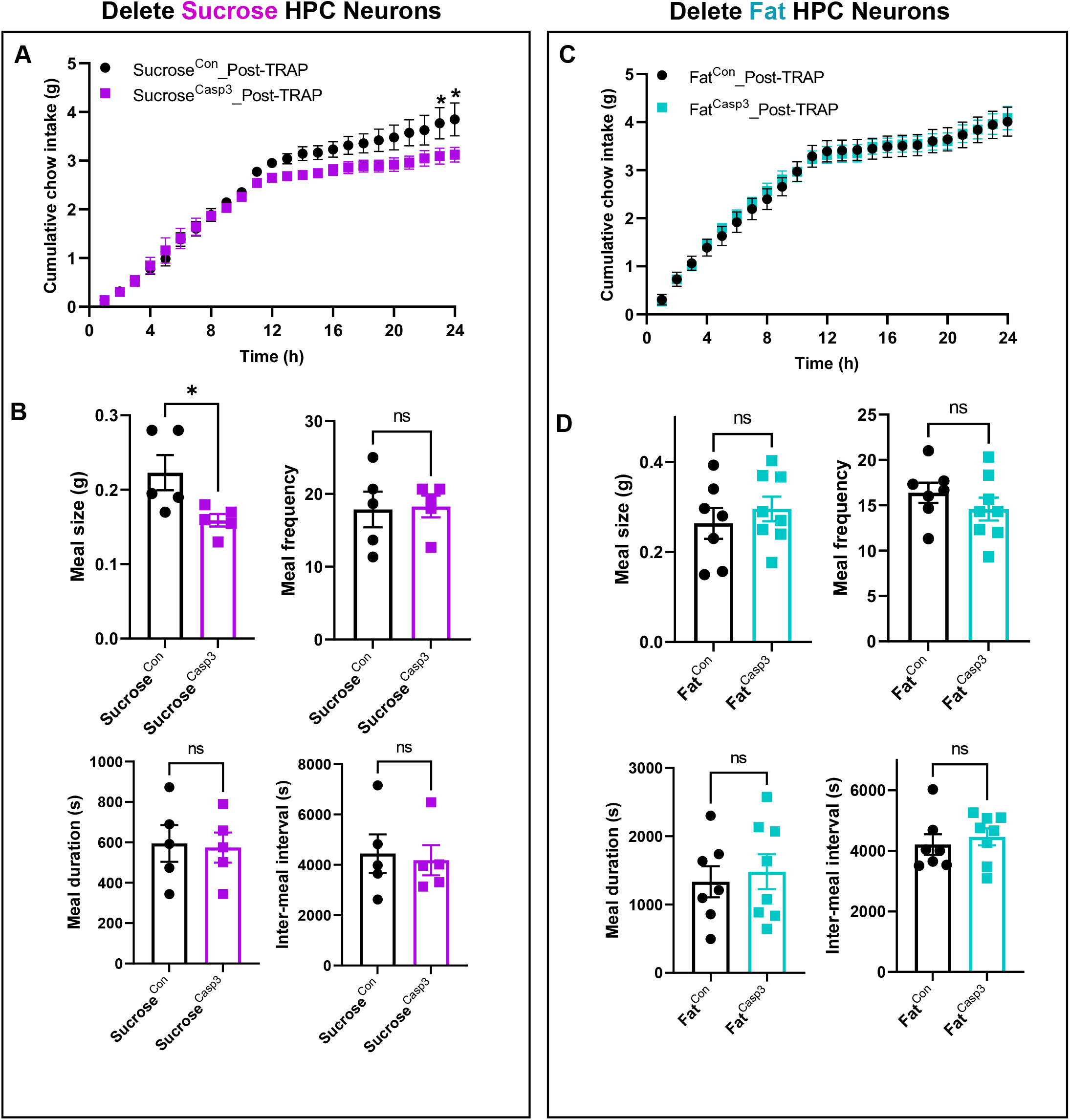
Sucrose-responsive dHPC neurons promote food intake. **A** Ablation of sucrose-responsive dHPC neurons reduces average daily chow intake, starting at dark onset (N=5/group, two-way ANOVA with Holm-Sidak post hoc analysis). **B** Ablation of sucrose-responsive dHPC neurons reduces meal size without affecting meal frequency, meal duration and inter-meal interval. (N=5/group, unpaired t test). **C** Ablation of fat-responsive dHPC neurons has no effect on chow intake (N=7-8/group, two-way ANOVA with post hoc analysis). **D** Ablation of fat-responsive dHPC neurons has no effect on meal pattern (N=7=8/group, unpaired t test). Data are presented as mean ± s.e.m. *P < 0.05, NS, not significant.

## Discussion

In the present study we identify novel populations of neurons in the HPC that influence decisions about where to locate food, what to eat, and how much to consume. We present multiple lines of evidence revealing distinct neural populations within the dHPC that respond to fat and sugar stimuli. Notably, these neurons respond to either fat or sugar infused into the gut, which allowed precise control over the volume and caloric content that each animal received, and isolated the post-ingestive effects from sensory cues like sight, smell and taste. These neuronal subsets not only exhibit spatial segregation in dHPC, but also exert distinct control of nutrient-specific preference and intake. Furthermore, they play a pivotal role in shaping feeding behavior through separate mechanisms involving memory, motivation, and preference.

### Identification of orexigenic neurons in the HPC

Prior studies have firmly established the crucial role of an intact HPC in the control of normal eating behavior. Notably, individuals with lesions that encompass the HPC diagnosed with anterograde amnesia have shown deficits in the regulation of satiety.^25, 76, 77^ In rodent models, pharmacological lesioning studies that remove the entire HPC have demonstrated an increase in food approach behavior,^26^ meal size^78^ and in some cases increased daily food intake and body weight.^11, 21, 22^ Furthermore, transient inhibition of neurons in either the ventral or dorsal HPC has been found to augment food intake,^23, 30, 31, 79, 80^ while stimulation of glutamatergic neurons decreases food intake.^23, 80^ Additionally, the administration of satiety hormones directly into the vHPC has been shown to decrease food intake, whereas the deletion of the receptors for these hormones increases food intake.^81, 82^ Together these data support the idea that the HPC plays a causal role in the inhibition of food intake. Yet the HPC is activated by ghrelin^34^ and food cues that both promote food intake,^36, 37^ and an orexigenic hippocampal circuit was recently discovered in humans and is strengthened in obesity.^40^ Circuits have been identified to connect HPC neurons with brain regions associated with motivated behavior,^23, 45, 83–89^ but these have for the most part not been linked with an increase in food intake. An exception is a vHPC to lateral hypothalamus circuit necessary to mediate endogenous ghrelin’s orexigenic effect in meal entrainment.^81^ Whether ghrelin receptor expressing vHPC neurons are an orexigenic population or if ghrelin inhibits an anorexigenic population remains to be determined. This study identifies two novel orexigenic populations in the dHPC.

We demonstrate a pivotal role of both dHPC neuronal populations in regulating the preference and intake of orally consumed solutions of isolated nutrients. Notably, deletion of nutrient-responsive dHPC neurons decreases intake and stimulation increases intake in a nutrient-specific manner. Specifically, dHPC^Fat^ neurons influence the quantity of fat solutions that animals consume, with no discernable impact on sugar intake. Conversely, dHPC^Sugar^ neurons exclusively govern sugar preference. Remarkably, our findings extend to complex diets. When exposed to chow, deletion of dHPC^Sugar^ neurons resulted in reduced food intake, whereas deletion of dHPC^Fat^ neurons had no effect. We found that dHPC^Sugar^ neurons are activated in response to both sucrose and starch, which suggests broad sensitivity to various carbohydrates. This could explain why chow - a grain-based diet composed of high levels of complex carbohydrate levels (63%, w/w), and low-fat content (<7%, w/w) – is impacted by dHPC^Sugar^, but not dHPC^Fat^ neurons. Collectively, our data suggest that the dHPC is highly attuned to the composition of the meal and separate recruitment of fat- or sugar-responsive dHPC neurons based on the nutrient composition can differentially influence food choice.

While previous studies have recognized a role for the HPC in responding to internal states of hunger and satiety,^12, 24–26^ the interoceptive HPC neurons responsible for this process had not previously been identified. We demonstrate that the vagus nerve is necessary for internal nutrient sensing in the gut to activate nutrient-responsive dHPC neurons that control food intake. Our findings build upon established evidence that connects the vagus nerve and the dHPC, including vagal-mediated HPC neurogenesis,^90–92^ synaptic function,^93, 94^ and the requirement of vagal sensory neurons that innervate the gut for optimal performance in hippocampal-dependent behavioral tasks.^47, 50^ Therefore, although the existence of a functional circuit was previously known, our data fills a gap in knowledge by pinpointing fats and sugars as physiological stimuli that activate this gut-hippocampal circuit to shape food preference.

### Explaining separate fat and sugar signaling mechanisms

One intriguing question that emerges from these findings is why distinct HPC neurons respond separately when activated by different post-ingestive stimuli? In the natural world, foods are rarely composed of a combination of both fat and sugar, potentially exerting selective evolutionary pressures that favored the development of separate biological systems for encoding distinct memories for either fat-rich or sugar-rich foods. Several studies have demonstrated that individuals exhibit more accurate spatial memory for the locations of high-calorie foods,^95–97^ hinting at the presence of memory systems finely tuned for efficiently locating and recalling nutritionally valuable food sources. These separate memory systems likely necessitate the ability to remember specific contextual cues associated with these different food types. We suggest that the formation of separate HPC neurons dedicated to either fat and sugar preference and appetitive memory may occur through one of two non-mutually exclusive mechanisms.

Firstly, ingested fats and sugars may activate separate parallel gut-brain circuits. Prior work from our lab provides evidence that fats and sugars are sensed by two separate populations of vagal sensory neurons.^46^ Notably, the deletion of these separate vagal populations was shown to impair learned preferences in a nutrient-specific manner.^46^ Furthermore, segregated cellular responses to fats or sugars in central reward circuits downstream of vagal sensory neurons were observed,^46^ suggesting the existence of separate hardwired signaling mechanisms for different nutrient reward. In support of this possible mechanism, we find that an intact vagus nerve is necessary for the response of dHPC neurons to either nutrient.

A second mechanism that would enable fat and sugar to activate separate dHPC populations is pattern separation. The DG in the HPC plays a pivotal role in the process of pattern separation,^98–101^ a fundamental computation that allows neural circuits to distinguish between similar input activity patterns and transform them into distinct output patterns.^102, 103^ This mechanism is crucial for avoiding the confusion of memories associated with similar experiences. Pattern separation has been well-established in rodent studies^104–109^ and is supported by human studies,^110, 111^ where the DG’s large number of neurons and sparse coding contribute to the decorrelation of input signals before reaching CA3.^103, 112^ Lesions to the DG result in novelty detection impairments following exposure to new spatial environments,^113^ highlighting its importance in reducing interference from previous experiences. We observe enrichment of DG activity in the dHPC in response to post-ingestive fat or sugar, which aligns with the possible role of the DG for discriminating contexts associated with appetitive compared to non-food related stimuli, but also encoding post-ingestive fat and sugar as dissimilar, non-overlapping memory representations.

### Identification of an appetitive engram for sugar

Neurons in the HPC play a pivotal role in transforming novel experiences into lasting memories that shape future behaviors. Immediate early genes (IEG), like Fos, are transiently expressed in specific HPC neuron populations following learned experiences.^114–116^ Reactivation of neurons based on IEG activity is essential for memory retrieval^117^ while inhibition of these ensembles impairs memory recall,^118^ underscoring the critical role of IEG in consolidating and recalling specific memories. The use of activity-dependent expression of reporters, therefore provides a framework for exploring engram ensemble. We utilize the Fos^TRAP^ mouse model to permanently tag activated neurons expressing Fos to target ensembles of appetitive stimuli. We find a network of Fos-expressing neurons in the dHPC responsive to the natural reinforcers, fat and sugar, that encode appetitive memory.

The term “engram” was originally introduced by Richard Semon to describe a memory representation.^119^ Since then, there have been ongoing efforts to locate the physical memory trace within the brain based on the ability to observe, erase and artificially express it as defining criteria.^120^ During the learning process, specific neuronal populations that constitute engram ensembles become activated and undergo cellular changes.^121, 122^ Inhibiting these changes impairs memory,^118^ while reactivation of these ensembles enable memory retrieval.^117^ Thus, significant progress has been made in understanding memory and engrams, particularly in the context of aversive and social interactions,^117, 123, 124^ but an engram associated with appetitive memory has not been defined, despite evidence that memory for food is highly conserved across species from insects to humans.^125–129^

We observe activation of a sparse population of neurons in response to post-ingestive nutrients, the first criteria of a memory trace. Selectively deleting sugar-responsive or fat-responsive neurons in the dHPC reduced nutrient-specific memory expression, satisfying the second criteria. Deleting the neurons tagged in response to sugar impaired the contextual memory for sugar, but had no effect on the subsequent expression of the contextual fat memory, and vice versa for fat responsive neurons. These findings support the idea that inhibiting components of one hippocampal engram does not affect expression of another separate engram^130^. Therefore, the sugar engram does not broadly disrupt memory retrieval, even of other appetitive memories.

Crucially, when we chemogenetically stimulated this specific ensemble of sugar-responsive dHPC neurons during training or testing, the mice exhibited improved performance in a highly-complex, contextually-dependent spatial memory task related to sugar. These findings strongly suggest that the dHPC^Sugar^ neurons contribute to the formation of a memory engram for sugar location and are sufficient for memory recall. Interestingly, activation of fat-responsive neurons did not improve performance in locating fat. Furthermore, ablation of the sugar-responsive neurons did not affect the learning a preference for a flavor associated with post-ingestive sugar, but led to rapid decline in the memory for the conditioned preference, highlighting the crucial role of these neurons in memory expression during the days following appetitive conditioning. Taken together, these data provide evidence of a sugar engram, and demonstrate that dHPC populations for fat and sugar are distinct.

In a separate set of experiments, we attempt to address the nature of the interrelationship between short-term memory (STM) and long-term memory (LTM). The debate revolves around whether these processes are distinct or part of a single memory system. Some argue that the STM system must be able to store complex representational structures that have never been encountered before,^131, 132^ while others propose a unified memory system^133–136^ or suggest that STM is an active component of LTM.^137–141^ Recent neuroimaging research has leaned towards the idea of a unified memory system,^134, 142–148^ although it may be difficult to parse out the overlapping features of STM and LTM that include encoding, retention, and recall. Our findings suggest that separate neural populations are involved in short-term working memory and long-term episodic memory. Specifically, deletion of sugar-responsive neurons in the dHPC impaired episodic memory, without impacting performance in a task of working memory.

### The hedonic hippocampus

The role of the hippocampus in motivated behavior remains unclear with mixed results from loss of function studies ^33, 149–151, 152^. In our study, we reveal that fat-responsive dHPC neurons are involved in both motivation and Pavlovian conditioning. When we deleted these neurons, mice displayed reduced effort to obtain fat reward, while stimulation increased their motivation to work for fat. These mice exhibited normal response during fixed ratio training, suggesting no impairment in learning. Furthermore, fat-responsive dHPC neurons are necessary and sufficient in flavor nutrient conditioning, providing clear evidence for a causal role for this small population of dHPC neurons in classical Pavlovian conditioning. Intriguingly, sugar-responsive neurons had no effect on motivation to work for sugar reward or formation of conditioned preference associated with sugar, aligning with previous studies indicating hippocampal lesions do not impact sugar conditioning.^73^ These results further underscore the different functions of fat and sugar dHPC neurons.

Gauthier and Tank (2018) identified a small but reliable population of hippocampal neurons that code for reward location across contingencies and environments.^64^ These neurons are thought to be involved in the process of encoding and retrieving memories related to rewards, regardless of the context in which the reward was experienced. The exact function of these reward anchored neurons has not yet been determined. Our data suggest a possible role for the reward location neurons for mapping the site of a reward and/or increasing the motivation to access the reward.

## Conclusion

The HPC is a brain region that is well known for its role in learning and memory, making it a candidate for supporting higher order decisions that underpin motivated behaviors, including feeding. Here we identify two novel populations of interoceptive dHPC neurons that are attuned to specific nutrients and allow highly-refined control over feeding behavior. We demonstrate that sugar-responsive dHPC neurons are part of an appetitive engram that encodes sugar location memory that can be erased or artificially activated. Conversely, fat-responsive dHPC neurons promote motivation and strengthen cue associations for post-ingestive fat. These neurons therefore have separate functions in creating an internal model that maps the environmental availability and locations of high-calorie foods (Sugar neurons) and modulates the internal drive to obtain them (Fat neurons). The role of these neurons in the pathophysiology of binge eating or obesity is yet to be explored. However, in our current food environment, there is the potential for devastating impact of these orexigenic neurons to exacerbate cue-induced consumption of obesogenic foods rich in fat and sugar.

## Supporting information

SupFig 1-5

## Acknowledgements

This study was supported by NIH grants R01 DK116004 (GL), R01 DK094871 (GL), R01 DK104897 (SEK), and T32 (MJM), in addition to start-up funds (GL), AHA postdoctoral fellowship (AS). The images used in figures 1A, 1F, 1K, 2A, 2I, 3A, 3J, 3G, 4A, 4F, 4I, S3B, and S3G were created using BioRender.com. We would like to acknowledge J. de Lartigue for constructive comments on the manuscript.

## Methods

### Animals and Housing

All animal procedures followed the ethical guidelines, and all protocols were approved by the Institutional Animal Care and Use Committee (IACUC) at the University of Florida (Protocol # 202110305) and Monell Chemical Senses Center (Protocol # 1187 and 1190). Adult mice (6-20 weeks of age of both males and females on a C57BL/6J background) were used and maintained on a reverse 12-h light/dark circle. Strain details and number of animals in each group are as follows: C57BL/6J wild type: n=48: 24 male, 24 female; bred in house by UF breeding core, Fos Cre Tomato: n=68: 34 male, 34 female; bred in-house from Jackson Laboratory B6.129(Cg)- Fos^tm1.1^(cre/ERT2)^Luo^/J (JAX stock no.021882) and Ai14 (B6.Cg-Gt(ROSA)26Sor^tm14(CAG-tdTomato)Hze^/J, JAX stock no.007914). Animals were single housed at 22°C with ad libitum access to standard rodent chow (3.1 kcal/g, Teklad 2018, Envigo, Sommerset, NJ) unless otherwise stated. We did not observe significant sex differences between male and female mice in our experiments. Prior to experiments, animals were habituated for 2-3 days to experimental conditions, including handling, injections, behavior chambers and attachment of gastric catheters for nutrient infusion.

### Surgeries

#### Vagotomy

Surgeries were performed aseptically following the IACUC Guidelines for Rodent survival surgery. Mice were anesthetized by inhalation of a continuous flow of 1.5-2% isoflurane. The pedal reflex test was performed prior to surgery to ensure that each mouse had reached an appropriate level of anesthesia. Mice were placed on a sterile drape warmed by a heating pad. Fur was shaved from the abdomen before cleansing with three exchanges of EtOH and Betadine. Sterile surgical equipment was used to create a 2-4cm midline laparotomy. The small intestine and colon were externalized and placed on sterile gauze moistened with sterile 0.9% NaCl saline. The subdiaphragmatic vagus nerve was visualized by gentle retraction of the liver and stomach. Complete vagotomy was performed by cutting the left and right cervical branches of the vagus directly caudal of the diaphragm using spring scissors. Sham animals had their subdiaphragmatic vagus nerve visualized, but not tampered with. The internal organs were repositioned and the incision site was covered with sterile gauze moistened with 0.9% NaCl saline until intestinal infusions.

Following the vagotomy, a silicone tubing was inserted via a small opening in the stomach wall, into the proximal section of the duodenal lumen. The duodenum received a 5-minute infusion of either sucrose (15%, w/v or fat (6.8%, v/v) solution (500 µL, 100 µL/min). Post-stimulation, incisions were sutured, and the mice were allowed to recover on a heating pad until they voluntarily moved to the unheated section of the cage. After 90 minutes, the mice were perfused and brains harvested, post-fixed in 4% PFA for 24 hours, and kept at 4 °C in a 30% sucrose in PBS solution until processing.

### Stereotaxic viral injections

Mice were anaesthetized with 1.5-2 % isoflurane and were injected with carprofen analgesia (5 mg/kg, s.c.) prior to bilateral injection in the dorsal hippocampus (dHPC). Core temperature was maintained using a homeothermic monitoring system and the absence of pedal reflex was utilized as a standard for appropriate depth of anesthesia. Animals were restrained in a stereotaxic frame (World Precision Instruments, Sarasota, FL) and their skulls were secured by positioning the bilateral ear crossbars into auditory meatus. A 2-3 mm incision was made in the midline of the scalp using a scalpel and the sagittal suture, bregma, and lambda of the skull were then exposed. With the bregma serving as an anatomical landmark, a dental drill was utilized to penetrate the skull above the target brain area. For dHPC viral injections, a Hamilton neuros syringe (Hamilton, Reno, NV) filled with a viral construct was lowered to the injection site in the dHPC (anteroposterior (AP): - 1.8 mm, mediolateral (ML): ± 0.4 mm, dorsoventral (DV): - 2.1 mm). The viral construct (0.2 µL/side, 0.1 µL/min) was injected via stereotaxic injector pump (Harvard Apparatus, Holliston, MA) and the needle remained in place for an additional 10 minutes to minimize the backflow of solution out of the injection site. The needle was removed slowly after the injection and 5-0 absorbable suture was used to close the skin. pAAV5-flex-taCasp3-TEVp was a gift from Nirao Shah and Jim Wells (Addgene viral prep # 45580-AAV5; http://n2t.net/addgene:45580; RRID:Addgene_45580),^59^ pAAV9-EF1a-DIO-hM3D(Gq)-mCherry was a gift from Bryan Roth (Addgene plasmid # 50460; http://n2t.net/addgene:50460; RRID:Addgene_50460), and pAAV9-EF1a-DIO-EYFP was a gift from Bryan Roth (Addgene viral prep # 44361-AAV9; http://n2t.net/addgene:44361; RRID:Addgene_44361).^153^

### Intragastric (IG) catheter implantation

IG catheters were made from 6 cm silicon tubing (.047” OD x .024” ID, SIL047, Braintree Scientific, MA) composed of 6 beads of silicon glue (#31003, Marineland, Blacksburg, VA) and a Pinport (Instech Labs, Plymouth Meeting, PA) for infusions. Analgesics buprenorphine XR (1 mg/kg) and carprofen (5 mg/kg) were injected (s.c.) 20 minutes prior to the surgery. Once animals had been anesthetized, a midline incision was made with a scalpel into the abdomen and hemostats were used to blunt dissect the skin layer away from the muscle layer to allow the catheter to be pulled between the abdominal incision site and the back of neck incision site. The stomach was exteriorized using a blunt forcep and a 4-mm purse suture was then placed at the junction of the greater curvature and fundus. Fine tip forceps were used to puncture the center of the purse suture and the end of the IG catheter was inserted into the stomach. The purse suture was then tightened and tied around the catheter. Next, a puncture hole was made in the left lateral abdominal wall using fine tip forceps and the catheter was pulled through and secured using 5-0 absorbable suture. The muscle layer of the abdominal incision site was then sutured closed and the open end of the catheter was pulled through to the back of the neck via a hole made in the middle of the shoulder blade. A 22-gauge Pinport was anchored in the tubing using superglue and once the patency of the catheter was confirmed via flushing with sterile saline, the catheter was secured with a purse suture around the hole in the back. Finally, the skin of the abdomen was closed with sterilized suture clips. For recovery, animals were fed with moistened chow in their home cage and were administrated Carprofen for 2 days after the surgery.

### Behavioral Tests

#### Food restriction

For all memory and motivation tasks involving food animals were maintained at 85-90% of their original body weight by food restriction. Briefly, for weight maintenance, the animals’ body weight was recorded every 24 hours and they were fed with a set amount of food calculated based on the loss of their original body weight. Animals were food restricted 6 hours before the task and not refed until 2h after the end of the task to prevent interference from food consumed outside of the task. If any mouse weighed less than 85% of their starting body weight, they were fed 2.5 g plus the excess weight loss until they reached 85% of starting body weight again. *Ad libitum* water access was provided in home cage.

### Food intake measurement

Food intake measurement and meal pattern analysis were performed using the BioDAQ episodic Food Intake Monitor (BioDAQ, Research Diets, Inc., New Brunswick, NJ). Previously validated meal criteria were used for food intake analysis (minimal meal size = 0.02 g, maximum inter-meal interval = 300 s).^154^ Animals were single housed and acclimated to the BioDAQ cages and fed ad libitum with chow for at least 3 days. Baseline food intake was then recorded for 3-5 days prior to performing the TRAP protocol, permitting within animal comparisons. Once the TRAP protocol was completed, animals were placed back in the BioDAQ cages, and their food intake was monitored for an additional 7 days. Meal parameters included meal size, the number of meals (meal frequency), meal duration and inter-meal interval were calculated by the BioDAQ Monitoring Software.

### Behavioral apparatus

Measurement of nutrient solution consumption and flavor-nutrient conditioning tests were conducted in mouse behavioral chambers enclosed in a ventilated and sound attenuating cubicle (Med Associates Inc., St. Albans, VT). Each chamber was equipped with slots for sipper tubing equipped with contact lickometers with 10 ms resolution (Med Associates Inc.) used for licking detection. All memory tests, except for the nutrient-driven Barnes Maze task, were conducted in open field apparatus (41x41 cm; 30 cm height). The foraging-related Barnes maze task^47^ involved an elevated white circular Barnes maze (Diameter: 92 cm, Height: 95 cm) with 16 holes (Diameter: 5 cm) evenly spaced around the outer edge of the table’s circumference. The holes were covered with petri dishes and visuospatial cues were placed on each of the walls surrounding the table. All memory tests were monitored and analyzed by tracking an animal’s head or body using the EthoVision XT Behavior Tracking Software.

### Nutrient solution consumption measurement

Food restricted mice were habituated and trained in these operant chambers with saccharin (0.2%, w/v) for 1 h/day for at least 3 days or until their total licking number reached at least 1,000 times/h. The bottle containing saccharin was placed in a different slot each day to avoid side preference. For caspase ablation studies, once animals were fully trained, they underwent consumption tests for either sucrose solution (15%, w/v) or isocaloric fat solution (6.8%, v/v) in a randomized order to minimize the influence of systematic contrast effects. Next, during consumption preference tests; one bottle with sucrose solution was placed on one side and another bottle with isocaloric fat solution placed on the other side. All the tests were conducted for 1 h/day for 3 days and the number of licks were recorded. For chemogenetic manipulation, baseline sucrose or fat consumption was assessed 20 min after saline injection. On the experimental day, sucrose or fat consumption was measured 20 min following the administration of clozapine-N-oxide (CNO; diluted in saline, 3 mg/kg, Enzo Life Sciences, NY).

### Two-bottle choice flavor nutrient conditioning test

To test whether ablation of nutrient-responsive HPC neurons affects specific nutrient-flavor association, a two-bottle preference test was performed. Once animals were considered trained to saccharin licking as described previously, a ‘pre’-test was performed in which they were given 10-min access to two novel Kool-Aid flavored solutions (cherry or grape, 0.05%, w/v) in saccharin (0.025%, w/v). To avoid side preference formation, sipper bottle positions were switched after 5 minutes. Subsequently, animals underwent a 1-hour conditioning session each day for 6 days where the lesser preferred flavor defined in the ‘pre’-test was paired with IG infusions of nutrients (CS+; 6.8% fat or 15% sucrose) and the preferred flavor was paired with IG infusions of saline (CS-). Specifically, during conditioning sessions, IG infusions of either nutrients or saline delivered by a syringe pump (20 µL/lick, 600 µL/min) were triggered by detection of the first lick and additional licks detected within 6 seconds had no programmed consequences. Upon completion of these conditioning sessions, mice underwent a ‘post’-test identical to the ‘pre’-test. The number of licks for the nutrient-paired flavor during ‘pre’ and ‘post’ tests was used to calculate flavor preference ratios (CS+ licks / total licks) before and after conditioning. For Sugar^TRAP^ mice, an additional ‘post’-test was also performed two days after the initial ‘post’-test to assess post-ingestive sucrose-conditioned flavor memory.

### Progressive ratio licking test

To assess whether nutrient-responsive dHPC neurons are important for nutrient-specific motivation, a progressive ratio (PR) operant licking test^71^ was performed. Food-restricted mice were initially trained to lick an active sipper spout to receive 15% sucrose or isocaloric fat solution via tubing mounted in a syringe pump (1 µL/lick, 600 µL/min) under fixed ratio (FR) 1 schedule (one hour/day for three days). After reaching >80% discrimination for the active sipper over the inactive sipper, the schedule was increased to FR5 for an additional two days. Tests under the PR schedule were then performed and failure to lick the active sipper in any 10 min period resulted in termination of the session (one hour/session). For chemogenetic experiments, on PR test days mice received either saline or CNO (i.p; 3 mg/kg) 20 minutes prior to entering the operant chamber followed by the opposite drug injection on the subsequent day. The number of licks was recorded and the breakpoint of reinforcement was calculated to quantify an animal’s willingness to work for a nutrient solution.

### Nutrient-driven food location memory test

To assess whether nutrient-responsive dHPC neurons are necessary for food location reference memory, a modified nutrient-driven food cup task^23, 155^ was conducted. Food restricted mice were habituated in open field apparatus, as described previously, for 5 minutes. The next day, a ‘pre’-test was performed where animals were allowed to explore the same arena containing two empty petri dishes placed in opposite corners for 5 minutes and the baseline preference for two quadrants was determined. Twenty-four hours later, animals underwent conditioning sessions (3 x 5 minute sessions) where the lesser preferred quadrant was paired with a petri dish containing drops of nutrient solution (CS+, 20 x 10 µL drops/session) and the preferred quadrant was paired with a petri dish containing drops of water (CS-, 20 x 10 µL drops/session). One or 24 hours after the last conditioning session, a ‘post’-test identical to the ‘pre’-test was conducted, i.e., both petri dishes were available with no stimuli. The time spent exploring each petri dish was recorded across the whole experiment and the discrimination index was calculated as the time spent exploring CS+ / total exploration time to assess animals’ memory performance.

### Nutrient-driven food location memory test with chemogenetic manipulation

To assess whether activation of nutrient-responsive dHPC neurons can improve context-dependent spatial memory for individual nutrients, a modified nutrient-driven food location memory test with chemogenetic manipulation with CNO was performed. Mice expressing hM3Dq or control virus in dHPC neurons trapped with intragastric infusion of sucrose (15% w/v) or fat (6.8% v/v) were habituated to two novel contexts (context A and context B). Two clean petri dishes were placed in opposing quadrants for each context. During a 3-day training phase, drops of either nutrient or water were added to the petri dishes (CS+, 20 x 10 µL drops/petri dish). Mice received saline injections 20 min prior to being placed in one context in the morning to learn to associate the location of a nutrient-containing dish for 10 min, and then received another saline injection 20 min before being placed in context B in the afternoon to learn a different location for the second nutrient-containing dish. Twenty-four hours after the last conditioning session, mice were tested with empty petri dishes in context A or context B to determine whether they could discriminate the correct context-specific nutrient-paired quadrant. After 7 days, the same test was repeated but CNO (3 mg/kg) was injected 20 min before mice were placed in the context which was paired with the nutrient with which the mice were trapped. To avoid desensitization, CNO was not injected on the third training day. The time spent exploring each petri dish was recorded across the whole experiment and the discrimination index was used to assess animals’ memory performance.

### Nutrient-driven Barnes maze test^47^

To evaluate the effect of nutrient-responsive dHPC neurons on food-related spatial working memory, a nutrient-driven Barnes maze test was performed. Food-restricted mice were first allowed to explore the Barnes maze apparatus for 5 min. The next day, animals were trained to utilize spatial cues to locate the correct petri dish containing sucrose (15% w/v) or isocaloric fat (6.8% v/v). All other petri dishes contained water. Each animal received two trials per day for three training days with 2-min inter-trial interval (during which time the maze is cleaned using 70% ethanol to avoid any confounding odor effect). Importantly, the target hole remains in a consistent position in both trials conducted on each training day but is relocated to a new position at the beginning of the initial trial on each subsequent training day. The number of incorrect investigations were recorded and the difference in the number of errors between trail 2 and trail 1 on an individual training day was calculated to determine whether animals improved their appetitive spatial working memory performance.

### Novel object in context (NOIC) test

To examine whether nutrient-responsive dHPC neurons only affect food-related memory, an HPC-dependent NOIC test^156^ was performed. Animals underwent 2 days of habituation (day 1 and day 2): half of the animals were allowed to freely explore context A, an opaque box with black stripes, for 10 minutes, whereas the other half were habituated to context B, an opaque box with no cues on the wall. The following day, groups were switched and habituated to the other context under the same environment. Training sessions were performed 24 hours after the last habituation session. On the training day (day 3), half of the animals were placed in context A for 10 minutes containing two identical blocks of Lego (object 1) placed in opposite corners, whereas the other half were first placed in context B for 10 minutes containing another two identical blocks of Lego (object 2) placed in opposite corners. Animals were placed back to their home cage between exposure to context A and context B and the inter-trial interval was 1-3 minutes. Animals were then switched and trained in the other context for another 10 minutes. On day 4 (test day), the NOIC recognition memory was tested by placing animals for 10 minutes in their last trained context (familiar context) containing one object (familiar object) which belonged to the familiar context on day 3 and one object (novel object) which belonged to the other context on day 3. The amount of time spent exploring each object was recorded and the discrimination index (DI) was calculated as [t_novel_/(t_novel_+t_familiar_)] in order to assess NOIC recognition memory. Seventy percent ethanol was used to clean all objects and contexts between tests.

## Histology

### TRAP protocol

As previously described,^46^ animals were fasted for 6 hours prior to IG infusion. Thirty minutes before onset of the dark phase, mice received an IG infusion of either sugar solution (15%, w/v) or fat solution (6.8%, v/v, Microlipid, Nestle, Vevey, Switzerland) (500 µL, 100 µL/min) in their home cage based on their assigned group. 4-hydroxytamoxifen (4-OHT, 30 mg/kg, i.p., MilliporeSigma, Burlington, MA) was injected 3 hours after the stimulus and standard chow was returned to animals’ home cage 3 hours after 4-OHT injection.

### Perfusions

Transcardial perfusion was performed in deeply anesthetized animals with phosphate buffer saline (PBS), followed by 4% paraformaldehyde (PFA). Following perfusion, brains were harvested and left in 4% paraformaldehyde for 24 hours and then transferred to a 30% sucrose solution containing 0.1% sodium azide for at least 72 hours before further processing.

### Tissue processing & storage

#### Slicing

Whole brains were frozen and embedded in OCT. A Leica frozen microtome (CM 3050 S, Leica Biosystems) was utilized to slice the frozen brains into 3 series at a thickness of 35 µm per section and slices were stored in cryoprotectant at -80°C until further staining or imaging.

#### Immunohistochemistry – Fos

The tissue was removed from cryoprotectant and rinsed in PBS 3 times (10 minutes/time) at room temperature. Subsequently, tissue was incubated for 30 minutes in a blocking buffer consisting of permeabilizing agent (244.5 mL of PBS, 5 mL of serum, 0.5 mL of Triton-X100, 0.25 g of Bovine Serum Albumin) and 20% normal donkey serum at 37 °C to prevent non-specific antibody binding. Tissue was then incubated overnight in PA containing a rabbit anti-cFos primary antibody (1:1000, Cell Signaling) at 4°C. On the following day, the tissue was rinsed in PBS 3 times for 20 each at room temperature followed by incubation in PA containing a donkey anti-rabbit IgG-AlexaFlour 647 secondary antibody (1:500, Abcam). Tissue was then rinsed in PBS 3 times for 1 hour each, mounted on slides, coverslipped with Prolong Diamond Antifade Mountant (Invitrogen, Waltham, MA), and stored at -20°C until imaging and analysis.

### Imaging

The HPC was identified using a mouse brain atlas (Paxinos and Franklin, 2001) and images of each region of interest were acquired with a Keyence BZ-X800 microscope using 10x objective. The number of positive cells, including trapped cells, cFos^+^ cells, and colocalization was counted automatically using NIS Element software with manual correction.

### Data analysis

Statistical analyses are described for each figure and were performed using GraphPad Prism 9 software. Two-tailed unpaired Student’s t tests were used for comparing two groups; Two-tailed paired Student’s t tests were used for comparing two treatments or tests in the same animal. One-way ANOVA, with or without repeated-measures, was used for comparing three groups; two-way ANOVA, with or without repeated-measures, was used for comparing more than one factor between groups. Data are presented as mean ± SEM and statistical significance is declared at *p* < 0.05.

## Supplemental Figures

**Supplemental Figure 1. vHPC neuronal response to intragastric nutrient infusions. A-B** Representative images of Fos expression in vHPC following intragastric infusion of saline, sucrose or fat. Scale Bar: (A) 100 µm, (B) 50 µm. **C-E** Quantification of Fos expression in dHPC and vHPC following intragastric infusion of saline, sucrose or fat. (N=3-5/group, paired Student’s t test). **F-G** Quantification of Fos expression in DG, CA3 and CA1 of the vHPC following intragastric sucrose or fat infusions (N=4-5/group, one-way ANOVA with Tukey post hoc analysis). **H** Quantification showing higher overlap between repeated infusions of fat compared to separate macronutrients in the dDG (N=5/group, unpaired t test). **I-J** Quantification of Fos^TRAP^ positive neurons and Fos immunofluorescence in the dHPC following intragastric sucrose or fat infusions, revealing a comparable density of responsive neurons (N=6/group, paired Student’s t test). **K** Representative images of tdTomato in the dHPC in response to sucrose (TRAP, magenta) compared with Fos-immunoreactivity following intragastric infusion of isocaloric starch (Fos, cyan) in the same animal. Scale Bar: 100 µm. **L** Quantification of (K) demonstrating that 60% of tdTomato labeled sucrose-responsive dHPC neurons co-express Fos activated by starch infusion (n=2/group). Data are presented as mean ± s.e.m. *P < 0.05, **P < 0.01, ***P < 0.001, **** P < 0.0001, NS, not significant.

**Supplemental Figure 2. In the absence of choice, fat-responsive dHPC neurons increase fat intake independently of taste. A-B** Ablation of sucrose or fat-responsive dHPC neurons has no effect on taste of sucrose or fat (N=8-9/group, unpaired t test). **C-D** Ablation of sucrose-responsive dHPC neurons has no effect on sucrose or fat consumption when individual solution is presented (N=8-9/group, two-way ANOVA with Holm-Sidak post hoc analysis). **E-F** Ablation of fat-responsive dHPC neurons decreases fat consumption without affecting sucrose consumption (N=8/group, two-way ANOVA with Holm-Sidak post hoc analysis). **G-H** CNO injection has no effect on sucrose or fat consumption in control virus-injected mice (N=5/group, two-way ANOVA with Holm-Sidak post hoc analysis). Data are presented as mean ± s.e.m. **P < 0.01, NS, not significant.

**Supplementary Figure 3. Nutrient-responsive dHPC neurons have no effect on nutrient-related working memory and non-food related memory. A** Animals exhibit a diminished ability to recall the nutrient-paired quadrant in the first trial, indicated by a reduced discrimination index (N=8-10/group, unpaired t test). **B** Schematic of novel object in context test. **C-F** Ablation of sucrose or fat-responsive dHPC neurons has no effect on non-food related memory test using novel object in context test (N=4-9/group, two-way ANOVA with Holm-Sidak post hoc analysis for C and E, unpaired t test for D and F). **G** Schematic of nutrient-related Barnes maze test. **H-O** Ablation of sucrose or fat-responsive dHPC neurons has no effect on sucrose-driven Barnes maze (H-K) and fat-driven Barnes maze test (L-O) (N=6-9/group, paired Student’s t test). Data are presented as mean ± s.e.m. *P < 0.05, **P < 0.01, NS, not significant.

**Supplemental Figure 4. Fat- and sucrose-responsive dHPC neurons are not required for learning. A** Ablation of sucrose-responsive dHPC neurons has no effect on the number of sucrose infusions acquired by animals during the sucrose conditioning sessions (N=4-6/group, two-way ANOVA with Holm-Sidak post hoc analysis). **B** Ablation of fat-responsive dHPC neurons reduces the number of fat infusions during the fat conditioning sessions (N=3-7/group, two-way ANOVA with Holm-Sidak post hoc analysis). **C-F** Sugar^TRAP^ and Fat^TRAP^ mice learned to lick in the active side to obtain infusion of sucrose or fat solution (N=3-7/group, two-way ANOVA with Holm-Sidak post hoc analysis). Data are presented as mean ± s.e.m. *P < 0.05.

**Supplemental Figure 5. Sucrose-responsive dHPC neurons control food intake. A** Ablation of sucrose-responsive dHPC neurons reduces chow intake compared to the baseline chow consumption within the same animal (N=5/group, two-way ANOVA with Holm-Sidak post hoc analysis). **B** TRAP protocol has no effect on chow intake in sucrose^TRAP^ mice expressing control virus (N=5/group, two-way ANOVA with Holm-Sidak post hoc analysis). **C** No group difference in baseline chow intake between Sucrose^Con^ and Sucrose^Casp3^ mice (N=5/group, two-way ANOVA with Holm-Sidak post hoc analysis). **D** Ablation of fat-responsive dHPC neurons has no effect on chow intake compared to baseline chow consumption within the same animal (N=3/group, two-way ANOVA with Holm-Sidak post hoc analysis). **E** TRAP protocol has no effect on chow intake in fat^TRAP^ mice expressing control virus (N=3/group, two-way ANOVA with Holm-Sidak post hoc analysis). **F** No group difference in baseline chow intake between Fat^Con^ and Fat^Casp3^ mice (N=3/group, two-way ANOVA with Holm-Sidak post hoc analysis).

