## Supplementary figures and images for "Separate orexigenic hippocampal ensembles shape dietary choice by enhancing contextual memory and motivation"

### SupFig 1-5

#### Supplemental Fig 1

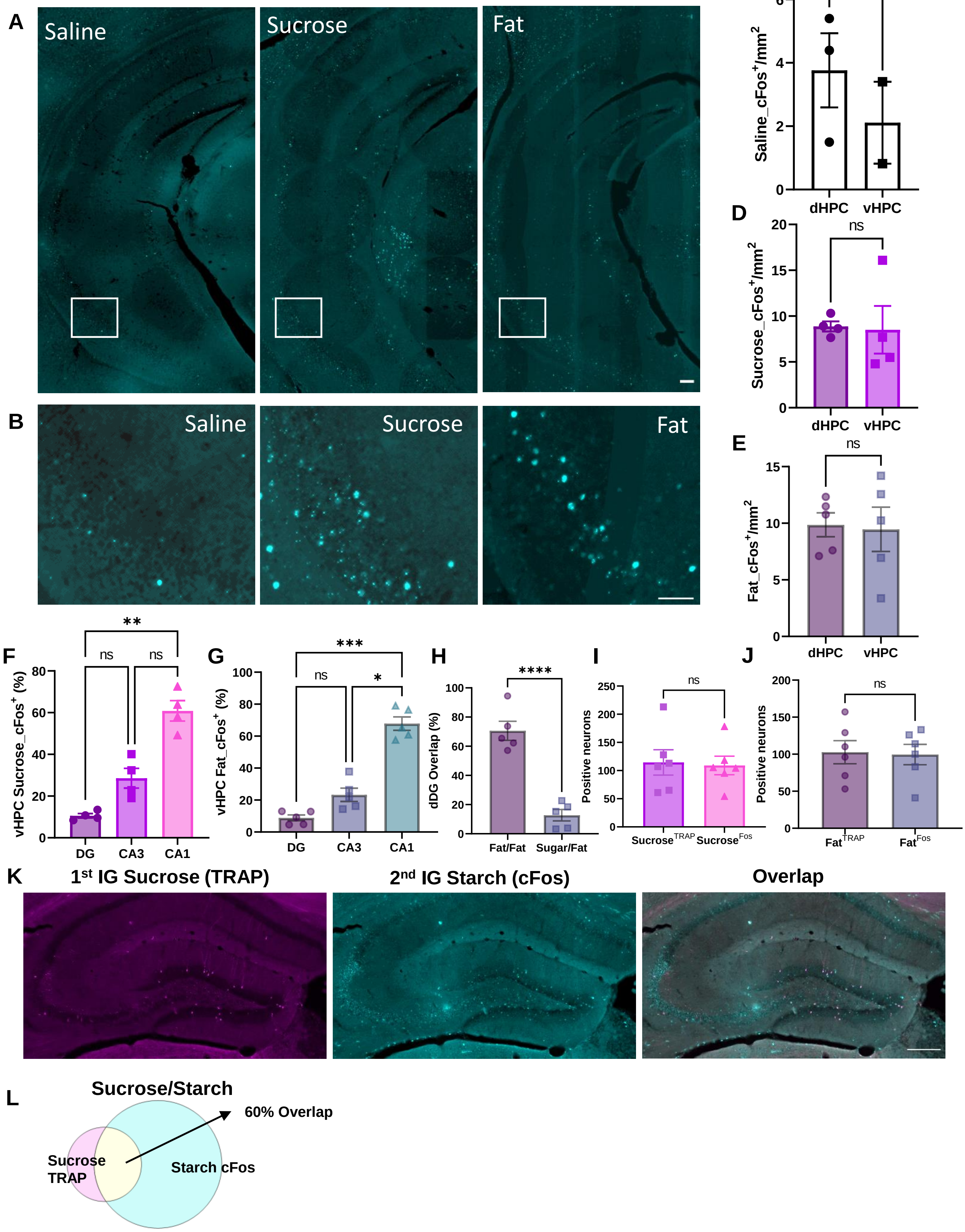

#### Supplemental Fig 2

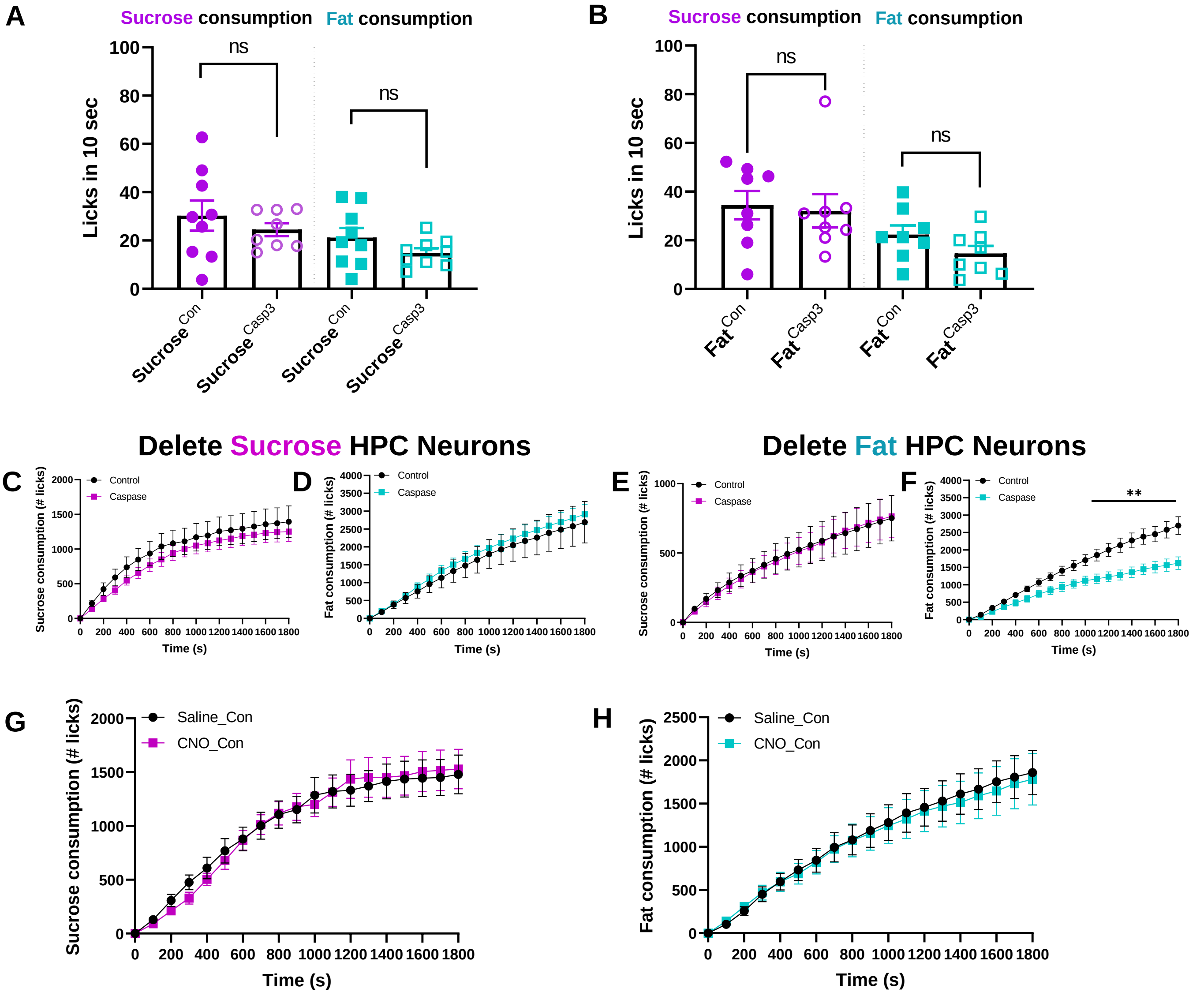

#### Supplemental Fig 3

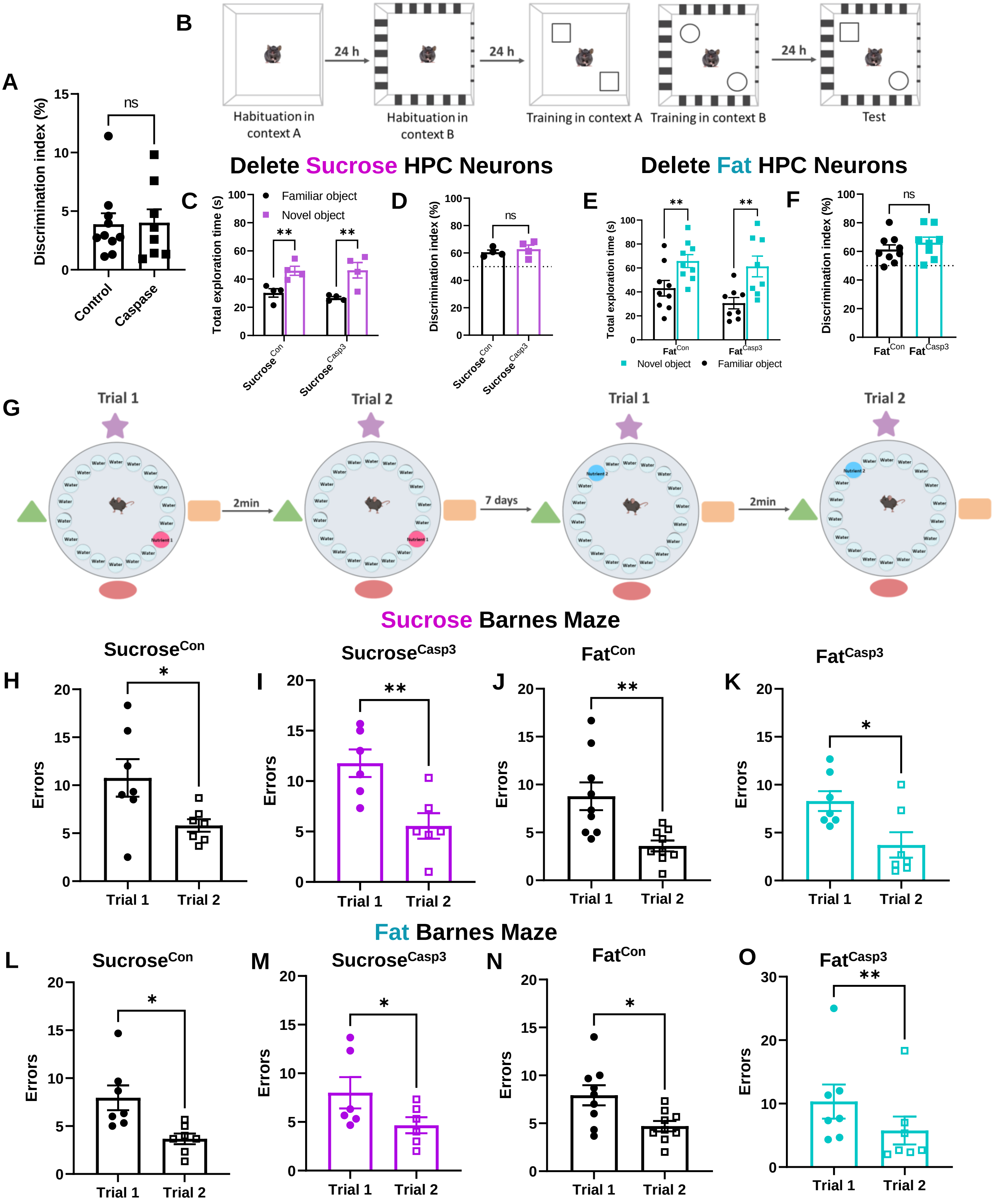

#### Supplemental Fig 4

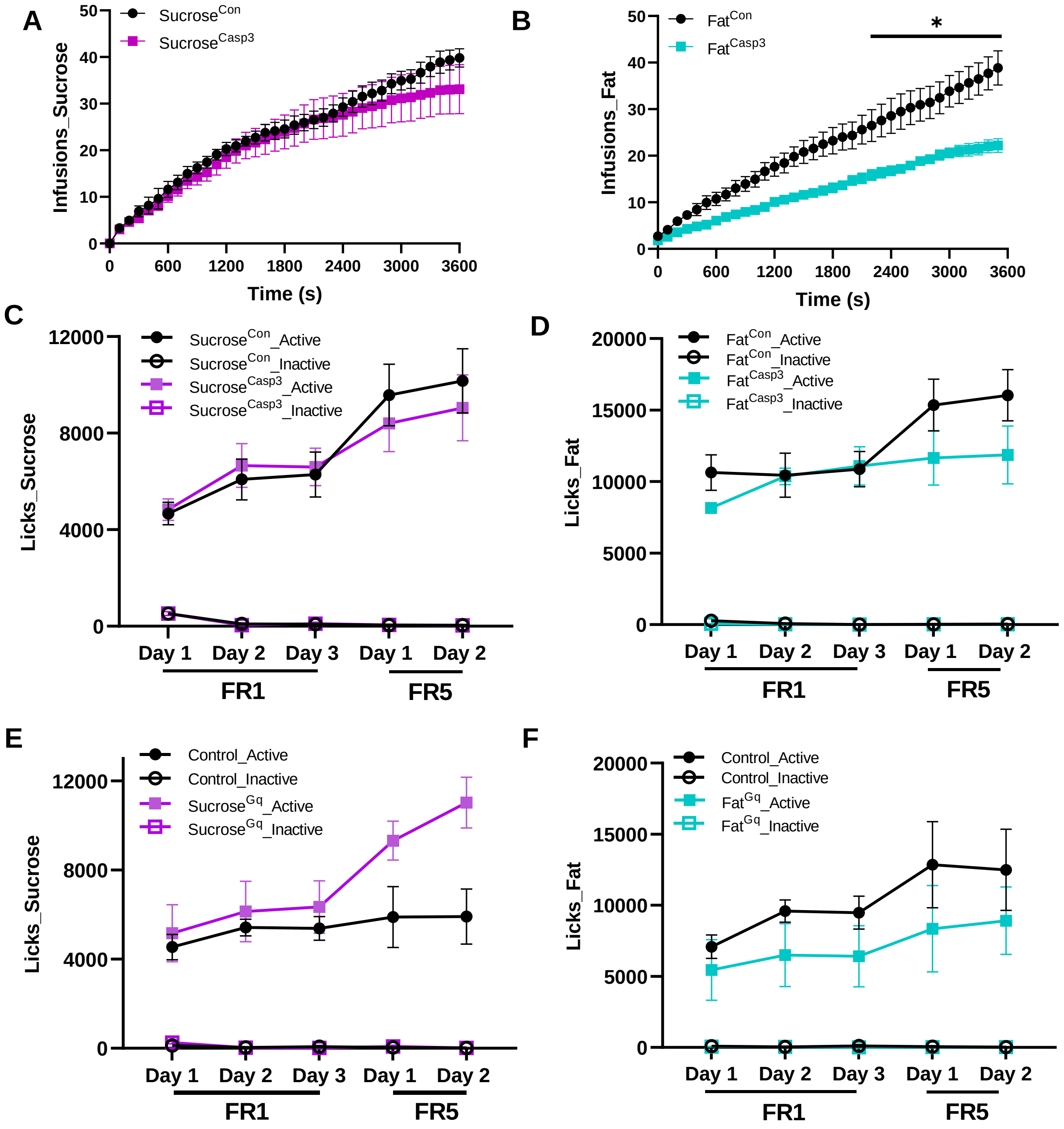

#### Supplemental Fig 5

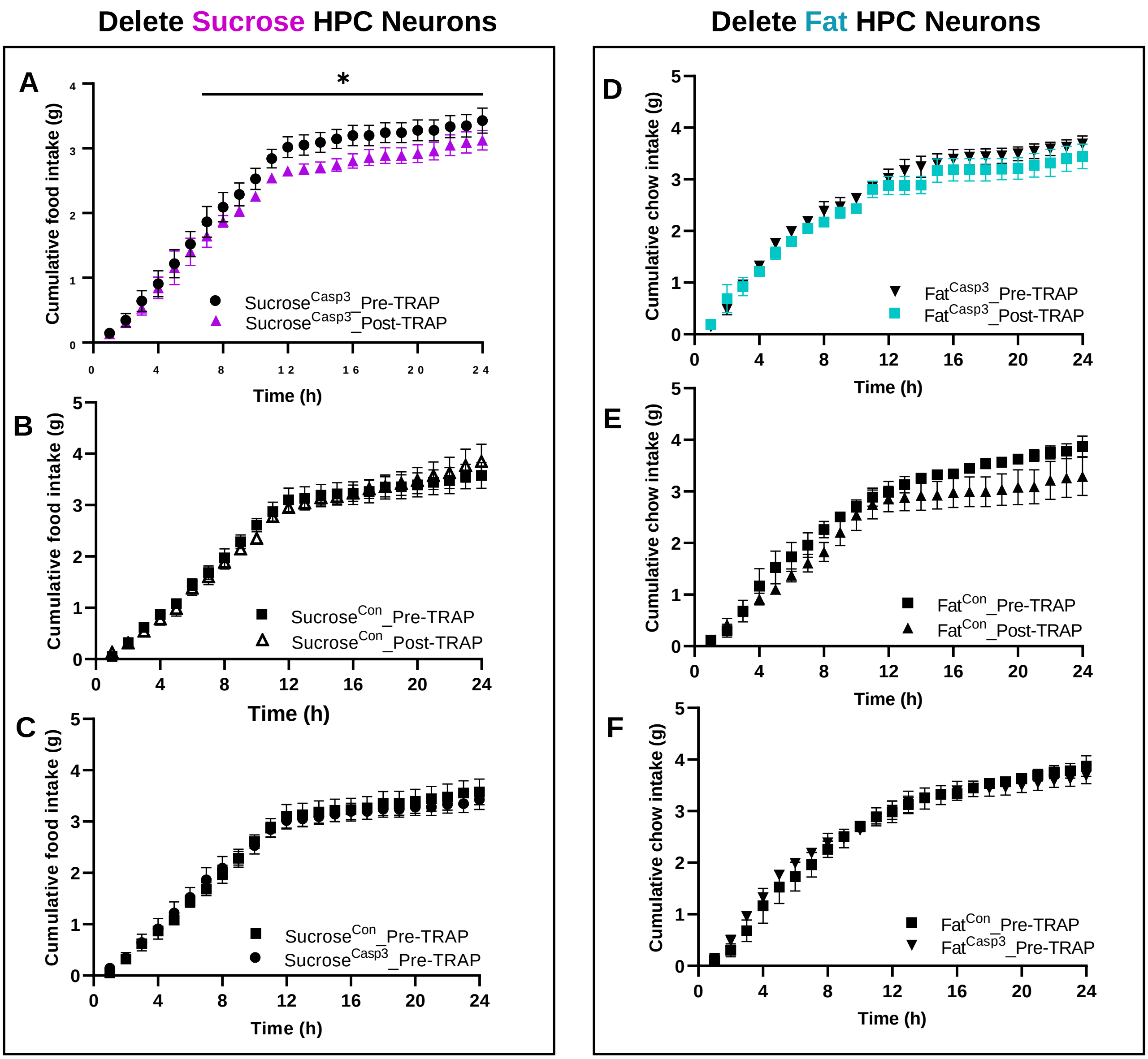
